# HB-EGF Signaling is Required for Glucose-Induced Pancreatic β-Cell Proliferation in Rats

**DOI:** 10.1101/683003

**Authors:** Hasna Maachi, Grace Fergusson, Melanie Ethier, Gabriel N. Brill, Liora S. Katz, Lee B. Honig, Mallikarjuna R. Metukuri, Donald K. Scott, Julien Ghislain, Vincent Poitout

## Abstract

The molecular mechanisms of β-cell compensation to metabolic stress are poorly understood. We previously observed that nutrient-induced β-cell proliferation in rats is dependent on Epidermal Growth Factor Receptor (EGFR) signaling. The aim of this study was to determine the role of the EGFR ligand Heparin-Binding EGF-like Growth Factor (HB-EGF) in the β-cell proliferative response to glucose, a β-cell mitogen and key regulator of β-cell mass in response to increased insulin demand. We show that exposure of isolated rat and human islets to HB-EGF stimulates β-cell proliferation. In rat islets, inhibition of EGFR or HB-EGF blocks the proliferative response not only to HB-EGF but also to glucose. Furthermore, knockdown of HB-EGF in rat islets blocks β-cell proliferation in response to glucose *ex vivo* and *in vivo* in transplanted glucose-infused rats. Mechanistically, we demonstrate that HB-EGF mRNA levels are increased in β cells in response to glucose in a Carbohydrate Response Element Binding Protein (ChREBP)-dependent manner. In addition, chromatin-immunoprecipitation studies identified ChREBP binding sites in proximity to the HB-EGF gene. Finally, inhibition of Src family kinases, known to be involved in HB-EGF processing, abrogated glucose-induced β-cell proliferation. Our findings identify a novel glucose/HB-EGF/EGFR axis implicated in β-cell compensation to increased metabolic demand.

## INTRODUCTION

In obesity, the maintenance of glucose homeostasis is dependent on the capacity of the pancreatic β cell to meet the increased insulin requirements that arise due to insulin resistance. Failure of this mechanism leads to type 2 diabetes (1). Hence, understanding how the β cell compensates for insulin resistance is a critical prerequisite to defining the pathogenesis of type 2 diabetes.

Β-cell compensation involves both an increase in the capacity to secrete insulin and an increase in mass. In adult rodents, β-cell expansion arises primarily from replication of existing β cells (2; 3). Over the last decade, modelling metabolic stress in rodents has led to the identification of an array of factors including insulin receptors (4), neurotransmitters (5), epidermal growth factor receptors (EGFR) (6), serpinB1 (7) and nutrients (8) that control β-cell proliferation. Prominent among these factors, glucose controls β-cell replication in rodent (9–12) and human (13) islets. Glucose-induced β-cell proliferation requires glucokinase, ATP-sensitive potassium channel closure and membrane depolarisation (10; 11). While several studies implicated insulin receptor signaling in glucose-induced β-cell replication (14; 15) this observation has been challenged by evidence supporting a role for Insulin Receptor Substrate 2 (IRS2), Mammalian Target of Rapamycin (mTOR) (16) and the Carbohydrate-Responsive Element Binding Protein (ChREBP) (17; 18). ChREBP is a glucose sensing transcription factor that binds DNA with its partner, Mlx, at carbohydrate response elements to stimulate glucose-responsive genes (19). Thus, the precise mechanisms underlying glucose-induced β-cell proliferation remains debated.

We established an *in vivo* model of nutrient excess in rats, in which a 72-h co-infusion of glucose and a lipid emulsion triggers a marked increase in β-cell proliferation and mass (20). Subsequent studies identified a signaling cascade involving the EGFR-mTOR-FoxM1 that underlies the β-cell response to nutrient infusion (21). In support of these finding, EGFR loss-of-function prevents compensatory β-cell mass expansion in adult rodents under conditions of physiological (pregnancy) and pathophysiological (high-fat feeding) insulin resistance (6) as well as following partial pancreatectomy (22). However, the identity of the EGFR ligand mediating this effect remains unknown. In previous studies we discovered that expression of the Heparin-Binding Epidermal Growth Factor (EGF)-like Growth Factor (HB-EGF) is up-regulated in islets from nutrient-infused rats, and that exogenous HB-EGF stimulates replication of MIN6 cells and primary rat β cells (21). HB-EGF is synthesized as a membrane-anchored precursor that is processed by the action of disintegrins and metalloproteinases (ADAM) to release the soluble active form (23). HB-EGF induces phosphorylation of EGFR and subsequent activation of a downstream signaling cascade including MAPK and PI3K/AKT.

The aim of this study was 1) to determine the role of HB-EGF in the β-cell proliferative response to glucose in rat islets *ex vivo* and *in vivo* and 2) to investigate the mechanisms linking glucose to an HB-EGF/EGFR signaling pathway promoting β-cell proliferation.

## RESEARCH DESIGN AND METHODS

### Reagents and solutions

RPMI-1640 and qualified FBS were from Invitrogen (Carlsbad, CA). Recombinant HB-EGF and betacellulin (BTC) were from R&D Systems (Minneapolis, MN). The HB-EGF inhibitor, CRM197, and the Src family kinase inhibitor, PP1, were from Sigma-Aldrich (St. Louis, MO). The EGFR tyrosine kinase inhibitor AG1478 and the mammalian target of rapamycin complex 1 (mTORC1) inhibitor rapamycin were from LC Laboratories (Woburn, MA). Adenoviruses expressing shRNAs against HB-EGF (Adv-shHBEGF) and control scrambled shRNA (Adv-shCTL) were from Vector Biolabs (Malvern, PA). SmartPool small interfering RNA (siRNA) duplexes against rat ChREBP and control siRNA were obtained from Dharmacon (Lafeyette, CO). Primary antibodies and dilutions are listed in Supplemental Table S1.

### Rat islet isolation and adenoviral infection

All procedures were approved by the Institutional Committee for the Protection of Animals at the CRCHUM. Islets were isolated from 2-month-old male Wistar or Lewis rats (Charles River, Saint-Constant, QC, Canada) by collagenase digestion and dextran density gradient centrifugation as described (24). For adenoviral infections, isolated islets were partially dissociated and then infected with 100 plaque-forming units of adenoviruses/cell overnight as described (25), after which the medium was replaced with complete medium and cultured for an additional 24 h prior to stimulation *ex vivo* or transplantation. To ensure that the HB-EGF knockdown was sustained for a period compatible with our *ex vivo* and *in vivo* experiments, we measured HB-EGF expression 5 days after infection. In Adv-shHBEGF-infected islets exposed to 16.7 mM glucose, HB-EGF mRNA was reduced by 32+/-8% (p<0.05; n=5) vs. Adv-shCTL-infected islets.

### Human islets

Islets from non-diabetic human donors were provided by the Alberta Diabetes Institute Islet Core and the Integrated Islet Distribution Program. The use of human islets was approved by the Institutional Ethics Committee of the CRCHUM (protocol no. ND-05-035).

### Islet proliferation *ex vivo*

Rat islets were cultured in RPMI-1640 with 10% (vol./vol.) qualified FBS (complete medium) for 72 h in the presence of glucose, 100 ng/ml HB-EGF or 50 ng/ml BTC as indicated in the Figure legends. EdU (10 μM) was added as indicated. The media were changed every 24 h. At the end of treatment, islets were embedded in optimal cutting temperature compound, frozen, sectioned at 8 μm and mounted on Superfrost Plus slides (Life Technologies Inc., Burlington, ON, Canada). Sections were immunostained for insulin (Ins) or Nkx6.1 to mark β cells, and for the proliferative markers Ki67, phospho histone H3 (pH3) or EdU (Click-iT™ EdU Imaging Kit, Life Technologies Inc.). Secondary antibodies were from Jackson ImmunoResearch (West Grove, PA). Images were acquired with a fluorescence microscope (Zeiss, Thornwood, NY). Proliferation was calculated as the percentage of double-positive Ki67+ (or pH3 + or EdU+) and Ins+ (or Nkx6.1+) cells over the total Ins+ (or Nkx6.1+) population. At least 1,500 β cells from 7-17 individual islets were manually counted per condition.

Human islets were hand-picked, washed with PBS and dispersed in accutase (Innovative Cell Technologies, Inc., San Diego, CA) for 10 min at 37°C. At the end of the digestion, cells were washed, resuspended and plated in 96-well plates (Perkin Elmer Inc., Waltham, MA) treated with Poly-D-Lysine Hydrobromide (Sigma-Aldrich, St. Louis, MO). After 24 h, dispersed human islets were cultured in RPMI-1640 with 1% (vol./vol.) human albumin serum (Celprogen, Torrance, Ca) for 72 h in the presence of glucose and 100 ng/ml HB-EGF as indicated in the Figure legends. The medium was changed every 24 h. At the end of treatment, cells were fixed and immunostained for insulin (Ins) and EdU. Images were acquired with an Operetta high-content imaging system (Perkin Elmer Inc., Waltham, MA) at 20x magnification. Approximately 1,500 cells were manually counted per condition.

### Static incubations

Triplicate batches of ten islets each were sequentially incubated twice with KRB solution containing 0.1% (wt/vol.) BSA and 2.8 mM glucose for 20 min at 37 °C, then incubated for 1 h with 2.8 or 16.7 mM glucose. Intracellular insulin content was measured following acid–alcohol extraction. Insulin was measured by radioimmunoassay using a rat insulin RIA kit (Millipore, Billerica, MA).

### Islet transplantation and glucose infusions in rats

Male Lewis rats weighing 250–350 g (∼2-month-old) (Charles River) underwent catheterization of the jugular vein for infusion and the carotid artery for sampling as described (26). For islet transplantation, 500 islets isolated from 2-month-old male Lewis rats were infected with Adv-shHBEGF or Adv-shCTL, as described above, and injected via a cannula under the left kidney capsule during the catheterization surgery. Animals were allowed to recover for 72 h followed by intravenous infusions of either saline (Sal) (0.9% wt/vol. NaCl; Baxter, Mississauga, ON, Canada) or 70% (wt/vol.) glucose (Glu) (McKesson, Montreal, QC, Canada) for an additional 72 h. The glucose infusion rate (GIR) was adjusted to maintain plasma glucose at 13.9–19.4 mmol/l throughout the 72-h infusion.

### Immunostaining of tissue sections

Transplanted kidneys and pancreata were fixed for 4 h in 4% paraformaldehyde and cryoprotected overnight in 30% sucrose. Tissues were then embedded in OCT, frozen, sectioned at 8 μm and mounted on Superfrost Plus slides (Life Technologies Inc.). Antigen retrieval was performed using sodium citrate buffer and β-cell proliferation was assessed as described above.

### Flow cytometry of β cells

Islets were isolated from male RIP7-RLuc-YFP transgenic rats (27), washed in PBS and dispersed in accutase for 10 min at 37°C. At the end of the digestion, cells were washed, resuspended in PBS, and passed through a 40-µm filter prior to sorting. Flow cytometric sorting of YFP-positive and -negative cells was carried out using a FACSAria II flow cytometer with FACSDiva software (BD Biosciences, San Jose, CA). YFP-expressing cells were detected using the 488-nm laser and 530/30-nm BP filter.

### Quantitative RT-PCR

Total RNA was extracted from 150-200 whole islets or 100,000 sorted islet cells using the RNeasy micro kit (Qiagen, Valencia, CA). RNA was quantified by spectrophotometry using a NanoDrop 2000 (Life Technologies Inc.) and 1 μg of RNA was reverse transcribed. Real-time PCR was performed by using QuantiTect SYBR Green PCR kit (Qiagen). Results were normalized to cyclophilin A RNA levels.

### Chromatin immunoprecipitation (ChIP) and chromatin confirmation capture (3C)

INS1 832/13 cell culture, siRNA treatment and RNA isolation and RT-PCR were performed as described (17). The pool of siRNA duplexes directed against ChREBP was previously shown to significantly decrease ChREBP mRNA (65%) and protein (70%) levels (17). ChIP was performed as previously described (18). Briefly, INS-1 cells were cultured for 16 h in 2 mM glucose followed by 6 h at 2 or 20 mM glucose. An anti-ChREBP or normal rabbit IgG were used for immunoprecipitation and a genomic region 30 kb downstream from the transcription start site of the HB-EGF gene known to bind ChREBP (28) was amplified by RT-PCR. 3C was performed essentially as described in (29). INS-1 cells were treated as for ChIP. The sequences of primers used for RT-PCR, ChIP and 3C are shown in Supplemental Table S2.

### Immunoblotting and ELISA

For immunoblotting, proteins were extracted from rat islets and subjected to 10% SDS-PAGE, transferred to nitrocellulose membranes, and immunoblotted with primary antibodies against phospho-EGFR, phospho-S6 ribosomal protein and α-tubulin in 5% (wt/vol) milk. Signals were revealed using horseradish peroxidase–conjugated anti-rabbit IgG secondary antibodies (Bio-Rad, Richmond, CA) in 5% (wt/vol) milk and visualized using Western Lighting Plus-ECL (Perkin Elmer Inc., Waltham, MA). Band intensity was quantified using ImageJ software (NIH).

HB-EGF was measured by ELISA (MyBiosource, San Diego, CA) in protein extracts from 200-300 rat islets treated with glucose for 1 h.

### Statistical analyses

Data are expressed as mean ± SEM. Significance was tested using one-way ANOVA with Tukey or Dunnett post hoc test, or two-way ANOVA with post hoc adjustment for multiple comparisons, as appropriate, using GraphPad Instat (GraphPad Software, San Diego, CA). P<0.05 was considered significant.

## RESULTS

### HB-EGF induces β-cell proliferation via EGFR-mTOR signaling

We previously showed that HB-EGF stimulates β-cell proliferation in dispersed rat islets (21). To confirm and extend these findings, we assessed the β-cell proliferative response to HB-EGF in intact rat islets after a 72-h exposure using either Ki67 or EdU-labeling to mark proliferating cells and insulin or Nkx6.1 to mark β cells (Fig. 1). In the presence of 2.8 mM glucose, 100 ng/ml HB-EGF or 50 ng/ml BTC increased the percentage of Ki67-positive β cells to levels comparable to those detected in response to 16.7 mM glucose (Fig. 1A-D). Similar results were obtained when using EdU as a proliferative marker (Fig. 1E&F). Exposing islets to the EGFR tyrosine kinase inhibitor AG1478 (300 nM) or the mTORC1 inhibitor rapamycin (10 nM) abrogated HB-EGF induced β-cell proliferation (Fig. 1G&H). In isolated dispersed human islets, exposure to HB-EGF for 72 h also induced β cell proliferation (Fig. 2).

**Figure 1.**
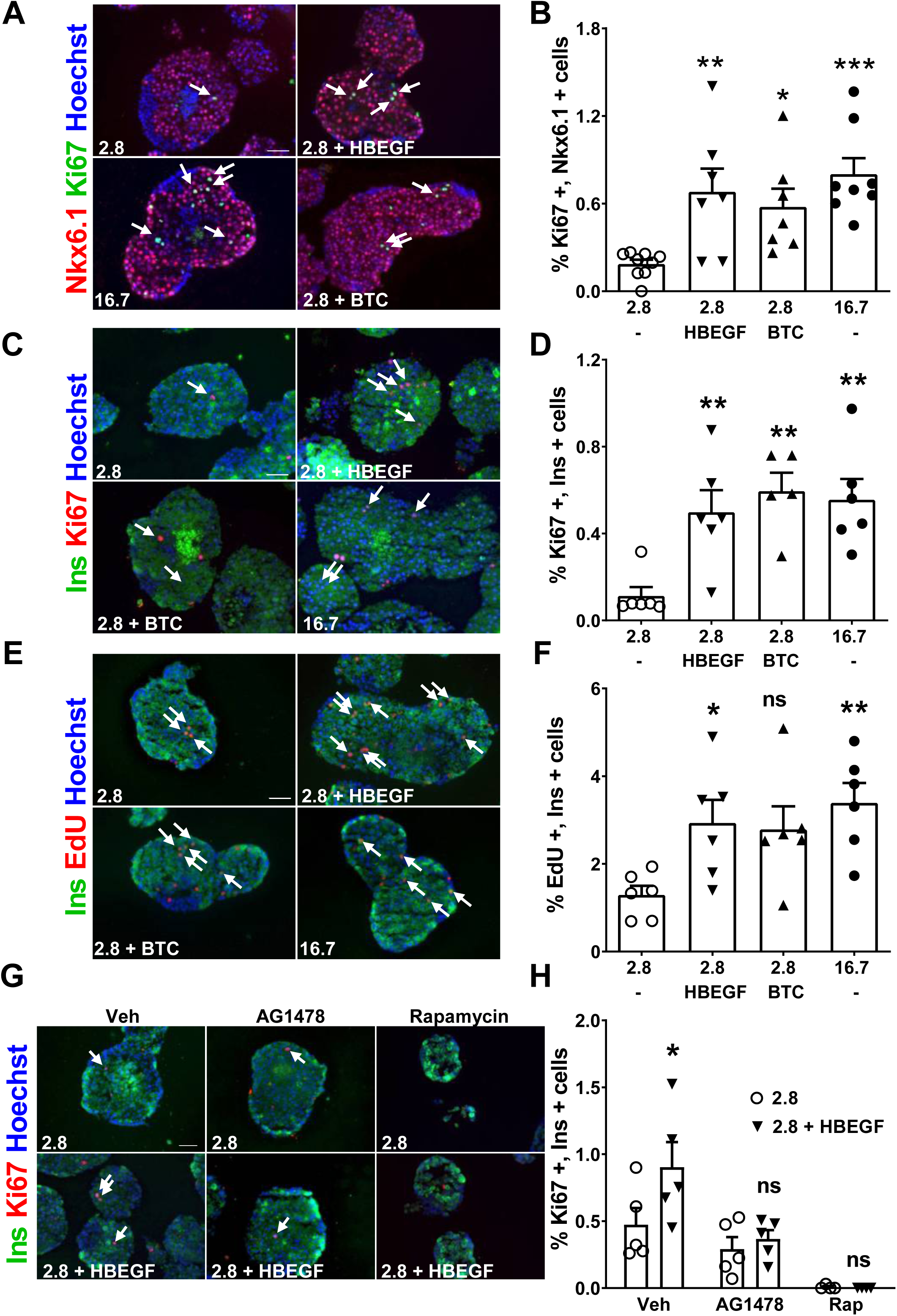
HB-EGF stimulates β-cell proliferation via the EGFR. (A-F) Isolated rat islets were exposed to 2.8 mM glucose, 16.7 mM glucose, or HB-EGF (100 ng/ml) or betacellulin (BTC; 50 ng/ml) in the presence of 2.8 mM glucose for 72 h. (G, H) Isolated rat islets were exposed to 2.8 mM glucose and left untreated or treated with HB-EGF (100 ng/ml) with or without AG1478 (300 nM) or rapamycin (Rap; 10 nM) for 72 h. Proliferation was assessed by Ki67 (A-D, G, H) or EdU (E, F) staining and Nkx6.1 (A, B) or insulin (Ins) (C-H). Representative images of Nkx6.1 (red), Ki67 (green) and nuclei (blue) (A) or insulin (green), Ki67 or EdU (red) and nuclei (blue) (C, E, G) staining. Arrows show positive nuclei for Ki67 and EdU. (B, D, H) The percentage of Ki67+ insulin+ (or Nkx6.1+) cells of total insulin+ (or Nkx6.1+) cells. (F) Percentage of EdU+ insulin+ cells of total insulin+ cells. Data represent individual values and are expressed as mean ± SEM (n=4–6). Scale bars, 50 µm. *p < 0.05, **p<0.01, ***p<0.001, as compared to the 2.8 mM glucose control condition. ns, not significant.

**Figure 2.**
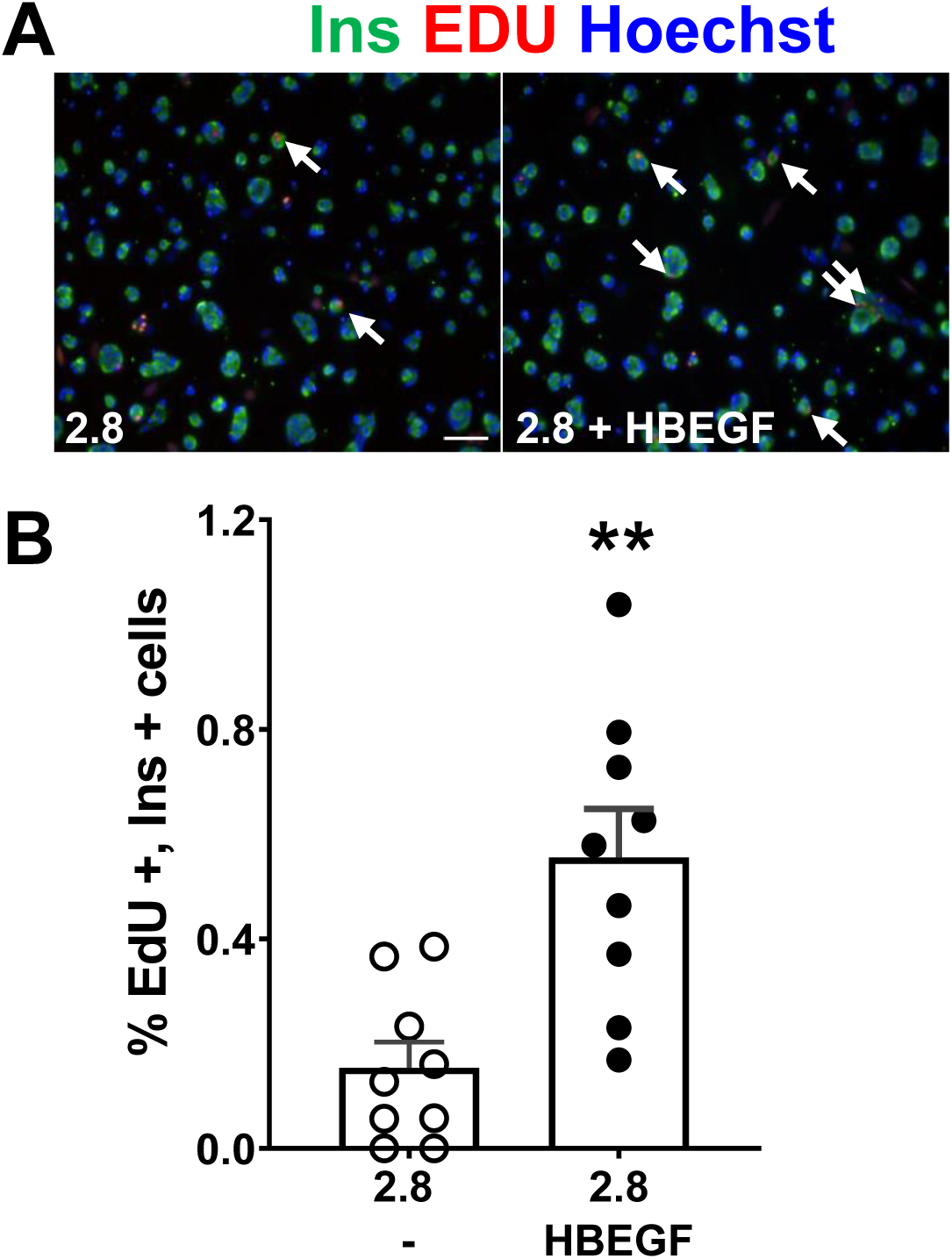
HB-EGF stimulates β-cell proliferation in human islets. Human islets were dispersed and then subjected to 2.8 or 16.7 mM glucose or HB-EGF (100 ng/ml) in the presence of 2.8 mM glucose for 72 h. Proliferation was assessed by EdU and insulin (Ins) to mark β cells. (A) Representative images of insulin (green), EdU (red) and nuclei (blue) staining. Arrows show positive nuclei for EdU. (B) The percentage of EdU+ insulin+ cells of total insulin+ cells. Data represent individual values and are expressed as mean ± SEM (n=9). Scale bars, 50 µm. **p<0.01, as compared to the 2.8 mM glucose control condition.

We then asked whether HB-EGF affects insulin secretion in rat islets. Isolated islets were either exposed to HB-EGF simultaneously with glucose during a 1-h static incubation to measure insulin secretion, or during the 24-h period preceding the static incubation. Neither acute nor prolonged exposure to HB-EGF significantly affected insulin secretion or insulin content (Supplemental Fig. S1). These results indicate that exogenous HB-EGF promotes rat β-cell proliferation via EGFR and mTOR without significantly affecting insulin secretion.

### Glucose-induced β-cell proliferation in isolated rat islets requires HB-EGF/EGFR signaling

Given that glucose is a known β-cell mitogen (9–12), we next examined the contribution of HB-EGF/EGFR signaling to glucose-induced β-cell proliferation. Treatment of rat islets for 72 h with 16.7 mM glucose led to an approximately 3-fold increase in Ki67 staining compared to 2.8 mM glucose (Fig. 3A&B). Addition of AG1478 completely prevented the glucose-induced increase in β-cell proliferation (Fig. 3A&B). Likewise, the HB-EGF inhibitor CRM197 (10 μg/ml) blocked the stimulatory effect of glucose on β-cell proliferation (Fig. 3C&D). Similar findings were obtained by labelling rat islets with Nkx6.1 and the M-phase marker pH3 after exposure to HB-EGF, 16.7 mM glucose, or 16.7 mM glucose + CRM197 (Fig. 3E&F). To further substantiate the implication of HB-EGF in glucose-induced β-cell proliferation, we infected isolated rat islets with Adv-shHBEGF or Adv-shCTL (Fig. 3G&H). Following a 72-h exposure to 16.7 mM glucose, Adv-shHBEGF-infected islets did not display any increase in β-cell proliferation (Fig. 3G&H). Collectively, these results demonstrate that HB-EGF/EGFR signaling is required for glucose-induced β-cell proliferation in isolated rat islets.

**Figure 3.**
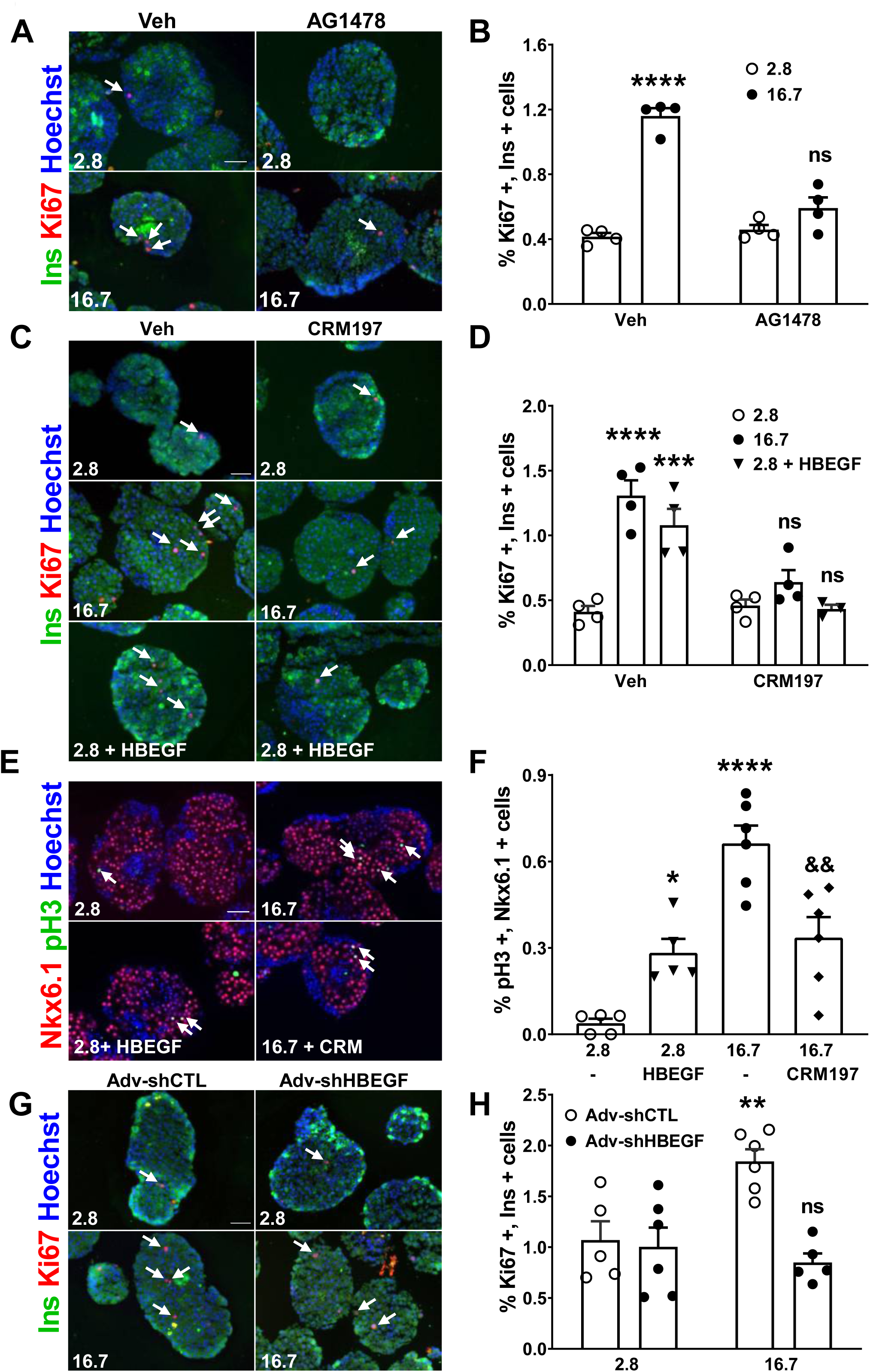
Glucose stimulates β-cell proliferation via HB-EGF/EGFR signaling. (A,B) Isolated rat islets were exposed to 2.8 or 16.7 mM glucose with or without AG1478 (300 nM) for 72 h. (C,D) Isolated rat islets were exposed to 2.8 mM glucose, 16.7 mM glucose, or 2.8 mM glucose + 100 ng/ml HB-EGF in the absence or presence of 10 μg/ml CRM197 for 72 h. (E,F) Isolated rat islets were exposed to 2.8 mM glucose with or without 100 ng/ml HB-EGF, or 16.7 mM glucose with or without 10 μg/ml CRM197, for 72 h. (G, H) Isolated rat islets were infected with Adv-shHBEGF or Adv-shCTL and exposed to 2.8 or 16.7 mM glucose for 72 h. Proliferation was assessed by Ki67 (A-D,G,H) or pH3 (E,F) staining and insulin (Ins; A-D,G,H) or Nkx6.1 (E,F) staining to mark β-cells. (A, C, E, G) Representative images of insulin (green) or Nkx6.1 (red), Ki67 (red) or pH3 (green) and nuclei (blue) staining. Arrows show positive nuclei for Ki67 or pH3. (B, D, H) Percentage of Ki67+ insulin+ cells of total insulin+ cells. (F) Percentage of pH3+ Nkx6.1+ cells of total Nkx6.1+ cells. Data represent individual values and means ± SEM (n=4– 6). Scale bars, 50 µm. **p<0.01, ***p<0.001, ****p<0.0001, as compared to the 2.8 mM glucose condition. && p<0.01, as compared to 16.7 mM glucose. ns, not significant.

### Glucose-induced β-cell proliferation in transplanted rat islets requires HB-EGF

To test whether islet-derived HB-EGF is necessary for glucose-induced β-cell proliferation *in vivo*, islets infected with either Adv-shHBEGF or Adv-shCTL were transplanted under the kidney capsule of Lewis rats. The rats were then infused with Sal or Glu for 72 h (Fig. 4A). Average blood glucose levels and glucose infusion rates were not different between both groups (Supplemental Fig. S2). As expected, the glucose infusion increased the percentage of Ki67-positive β cells in the endogenous pancreas to the same extent in Adv-shCTL and Adv-shHBEGF transplant recipients (Fig. 4B&C). Adv-shCTL-infected islet grafts also showed increased β-cell proliferation in response to glucose infusion (Fig. 4D&E). In contrast, Adv-shHBEGF-infected islets were unresponsive to glucose (Fig. 4D&E). These data demonstrate that, as observed in isolated islets (Fig. 3), HB-EGF/EGFR signaling is required for glucose-induced β-cell proliferation *in vivo*.

**Figure 4.**
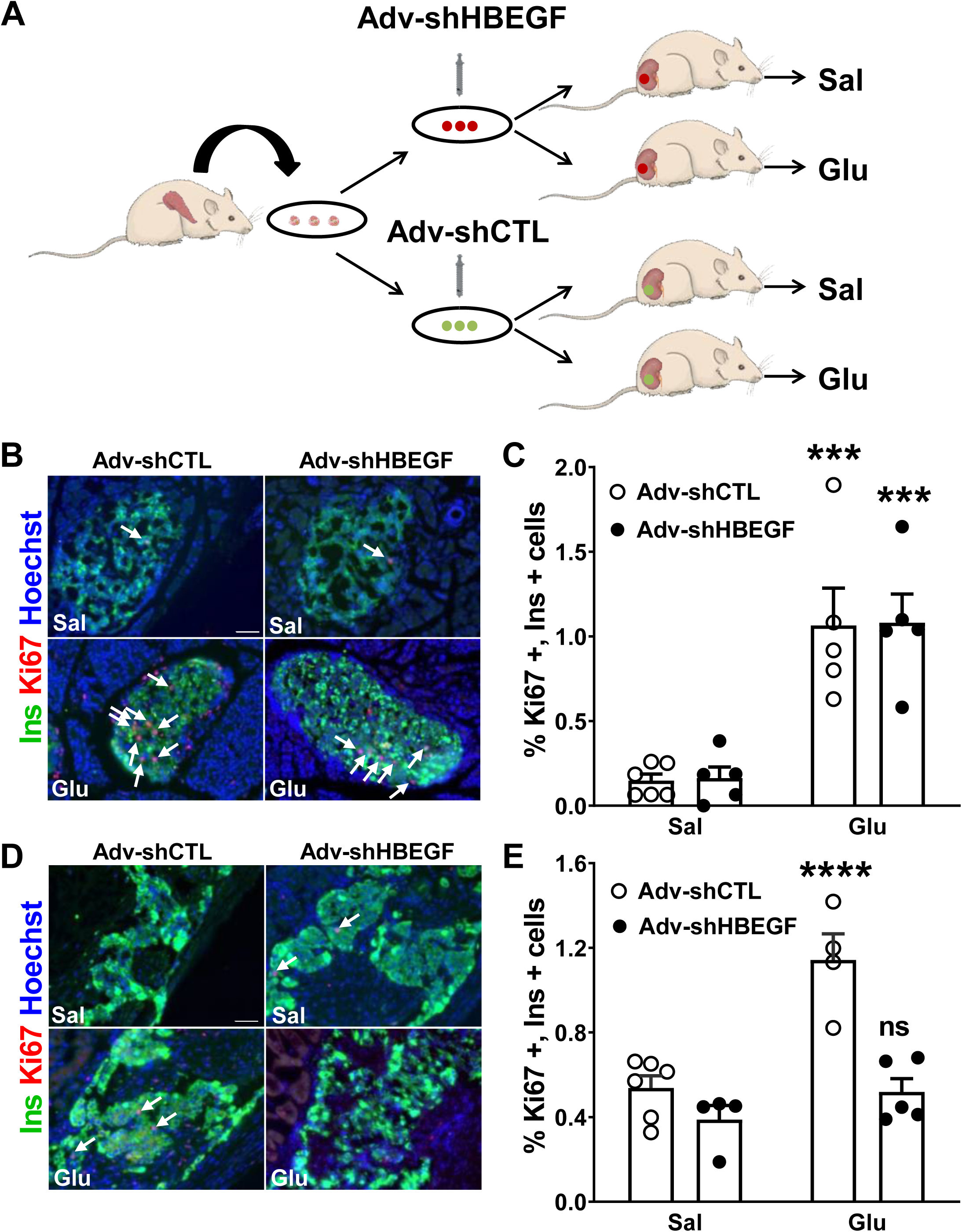
HB-EGF is required for glucose-induced β-cell proliferation *in vivo*. (A) Isolated rat islets were infected with Adv-shHBEGF or Adv-shCTL and transplanted under the kidney capsule of 2-mo-old Lewis rats infused with saline (Sal) or glucose (Glu) for 72 h. (B-E) Proliferation was assessed by Ki67 staining and insulin (Ins) staining. (B, D) Representative images of insulin (green), Ki67 (red) and nuclei (blue) staining in the pancreas (B) or transplanted islets (D). Arrows show positive nuclei for Ki67. (C, E) The percentage of Ki67+ insulin+ cells of total insulin+ cells in the pancreas (C) and transplanted islets (E). Data represent individual values and means ± SEM (n=4–6). Scale bars, 50 µm. ***p<0.001, ****p<0.0001, as compared to the saline condition. ns, not significant.

### HB-EGF gene expression is up-regulated in β cells in response to glucose

As we previously showed that infusion of glucose and lipids in rats increases HB-EGF mRNA levels in islets (21), we asked whether glucose alone was sufficient to stimulate HB-EGF expression in isolated islets. Indeed, isolated rat islets exposed to 16.7 mM glucose for 24 h displayed a 1.5-fold increase of HB-EGF mRNA compared to 2.8 mM glucose (Fig. 5A). To determine whether the increase in islet HB-EGF gene expression was primarily in β cells, we used a transgenic rat expressing yellow fluorescent protein (YFP) under the control of the Ins2 promoter (RIP7-RLuc-YFP) (27) to enrich for β cells by flow cytometry after glucose treatment. Glucose augmented HB-EGF mRNA levels in the YFP-positive (β-cell enriched; Fig. 5B) cells, but not the YFP-negative (Fig. 5C) fraction, suggesting that glucose stimulates HB-EGF gene expression in rat β cells.

**Figure 5.**
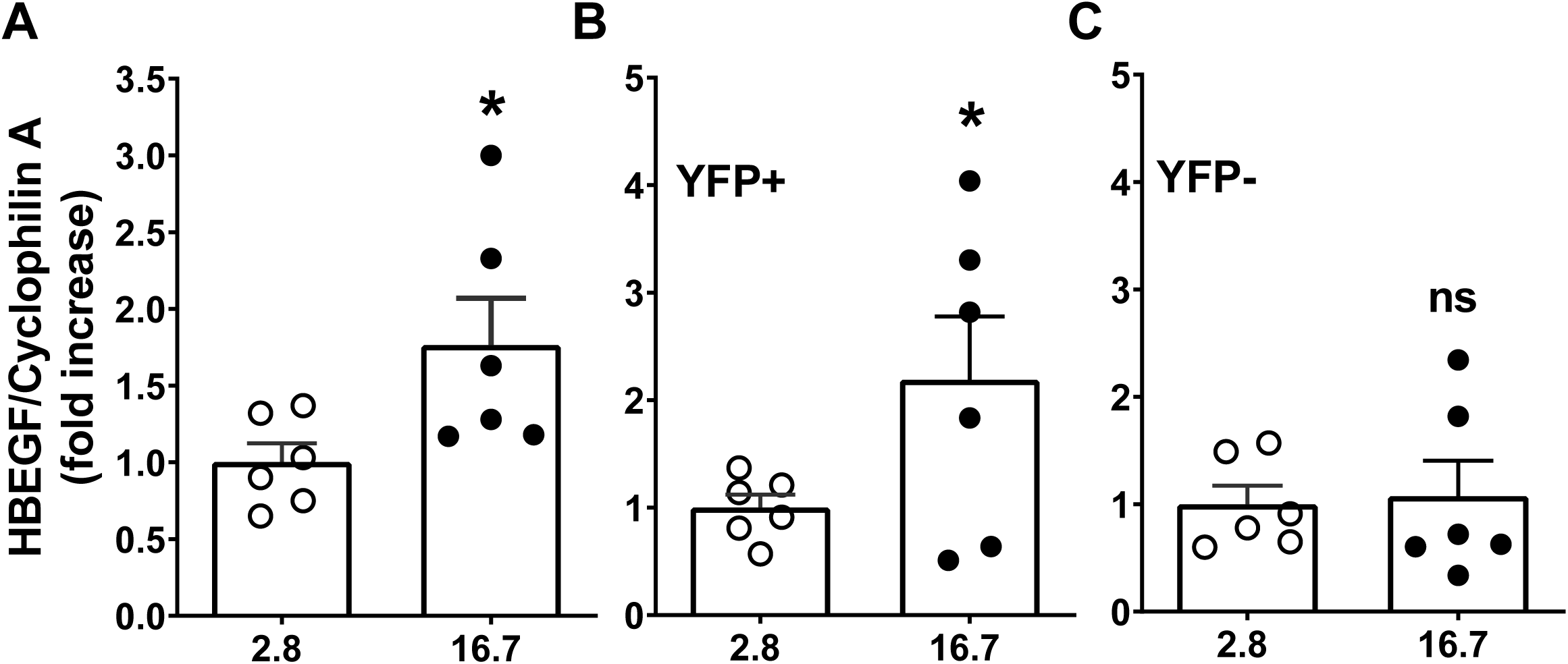
Glucose increases HB-EGF gene expression in the β-cell. (A-C) HB-EGF mRNA was measured in isolated intact rat islets (A) or in FACS-sorted YFP-positive (B) and YFP-negative (C) cells from isolated RIP7-RLuc-YFP islets following exposed to 2.8 or 16.7 mM glucose for 24 h. mRNA was determined by quantitative RT-PCR and normalized to cyclophilin A. Data are presented as the fold-increase over the 2.8 mM glucose condition and represent individual values and means ± SEM (n=5-6). *p < 0.05, as compared to the 2.8 mM glucose condition. ns, not significant.

### Glucose stimulates HB-EGF gene expression via ChREBP

ChREBP is a key mediator of glucose-induced transcriptional changes (28). Therefore, we asked whether HB-EGF is a direct target of ChREBP. Consistent with the results shown in Fig. 5, glucose increased HB-EGF expression in untransfected INS-1 cells and in cells transfected with a control siRNA (Fig. 6A). In contrast, siRNA-mediated knockdown of ChREBP abolished the glucose response (Fig. 6A). ChREBP ChIP-seq and DNase-seq analyses of INS-1 cells exposed to glucose identified putative enhancer elements containing canonical ChREBP binding sites located approximately 30 kb downstream of the HB-EGF transcriptional start site (28). ChIP analysis for one of these elements showed that a 6-h exposure to 20 mM glucose significantly increased ChREBP binding, whereas binding to a control region was unchanged (Fig. 6B). Furthermore, 3C analysis revealed increased interactions between these enhancers and the HB-EGF promoter in the presence of 20 mM glucose (Fig. 6C). These results show that glucose-induced HB-EGF gene expression is mediated by direct binding of ChREBP to enhancers located 3’ to the HB-EGF gene.

**Figure 6.**
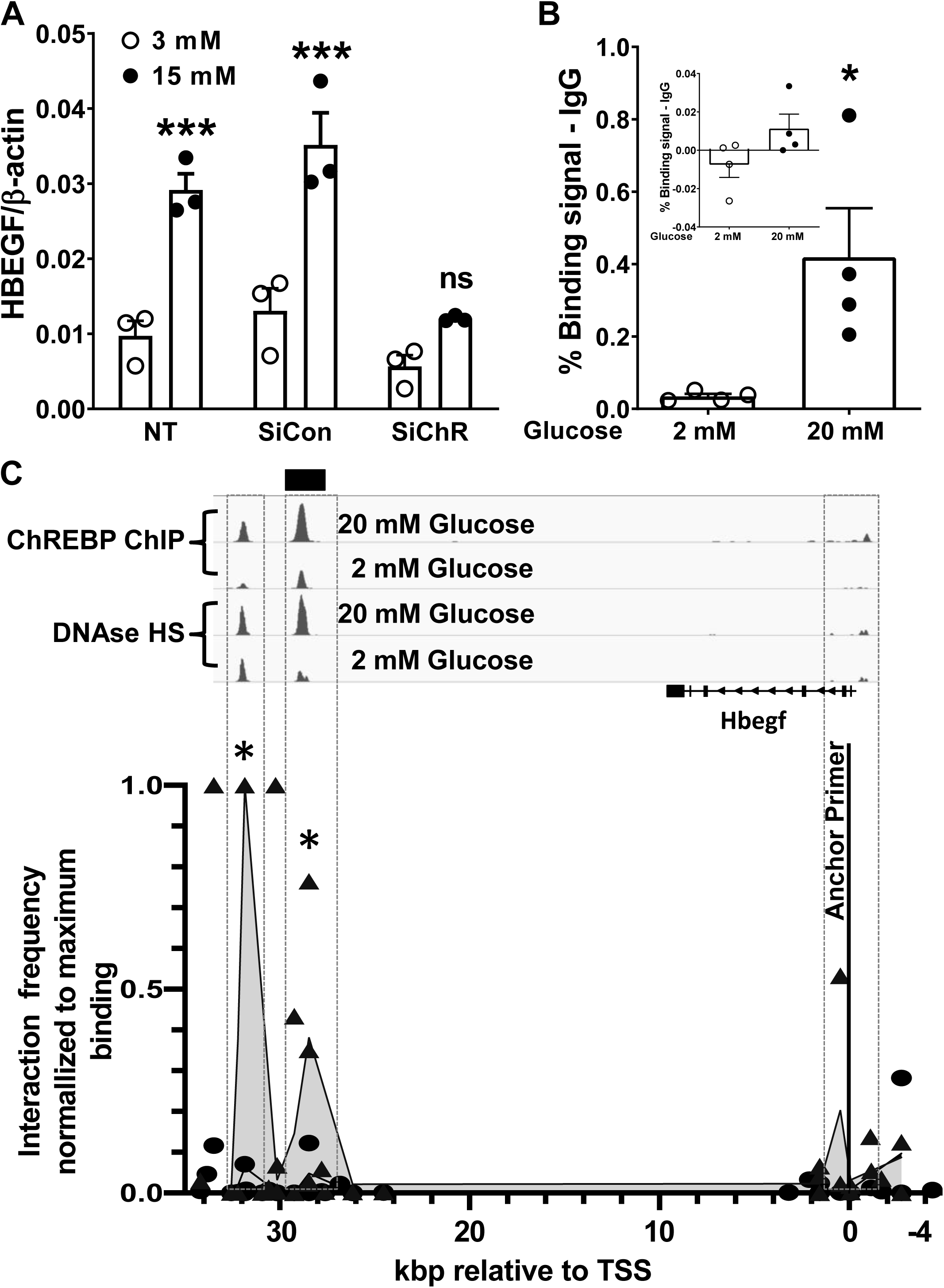
ChREBP mediates glucose-induced HB-EGF gene expression in INS-1 cells. (A) HB-EGF RNA was measured in INS-1 cells exposed to 3 or 15 mM glucose for 18 h in the presence of a control siRNA (SiCon) or an siRNA directed against ChREBP (SiChR) (n=3). mRNA was determined by quantitative RT-PCR and normalized to β-actin. NT, non transfected. (B) ChREBP binding to a genomic region 30 kb downstream from the transcription start site of the HB-EGF gene known to bind ChREBP (black bar in C, upper) was assessed in INS-1 cells exposed to 2 or 20 mM glucose for 6 h followed by ChIP using an antibody against ChREBP or control IgG (n=3). Data indicate the percent binding after subtraction of the IgG control. Inset, Pklr coding region serves as a negative control. (C) Upper, genome browser view of 38,000 bp of the genomic locus spanning the TSS of the HB-EGF gene showing the ChREBP binding (ChREBP ChIP) and DNAse hypersensitivity sites (DNAse HS) downstream of the gene (28). Black bar, region amplified in (B). Lower, chromatin confirmation capture (3C) data from INS-1 cells treated as in (B), aligned to the genome browser and expressed as interaction frequency normalized to maximum interaction (n=3). Black line, anchor primer. Shaded gray added for clarity represents interaction frequency after 20 mM glucose treatment. Data are expressed as mean ± SEM.*p < 0.05, ***p<0.001, as compared to the control condition. ns, not significant.

### Glucose-induced β-cell proliferation is dependent on Src upstream of EGFR activation, but glucose-induced mTOR activation does not require HB-EGF

Processing of proHB-EGF by ADAM proteins releases the active form that binds and activates EGFR (23). Previous studies in mesangial cells suggest that glucose-induced proteolytic processing of HB-EGF requires Src activation (30). Therefore, we investigated the role of Src family kinases in glucose-induced β-cell proliferation. Addition of the Src inhibitor PP1 abrogated the β-cell proliferative response to 16.7 mM glucose but not to HB-EGF (Fig. 7A&B), consistent with the possibility that glucose promotes proHB-EGF cleavage via Src followed by HB-EGF activation of EGFR. To assess glucose-stimulated HB-EGF shedding, we attempted to measure HB-EGF levels in islet conditioned media following a 1-h exposure to 16.7 mM glucose. Unfortunately, HB-EGF levels in the samples were below the detection limit of the assay. However, we observed a trend towards an increase in total HB-EGF levels in islet extracts (Supplemental Fig. S3) which, although not statistically significant, is consistent with the glucose-induced HB-EGF expression shown in Fig. 5A & 6A.

**Figure 7.**
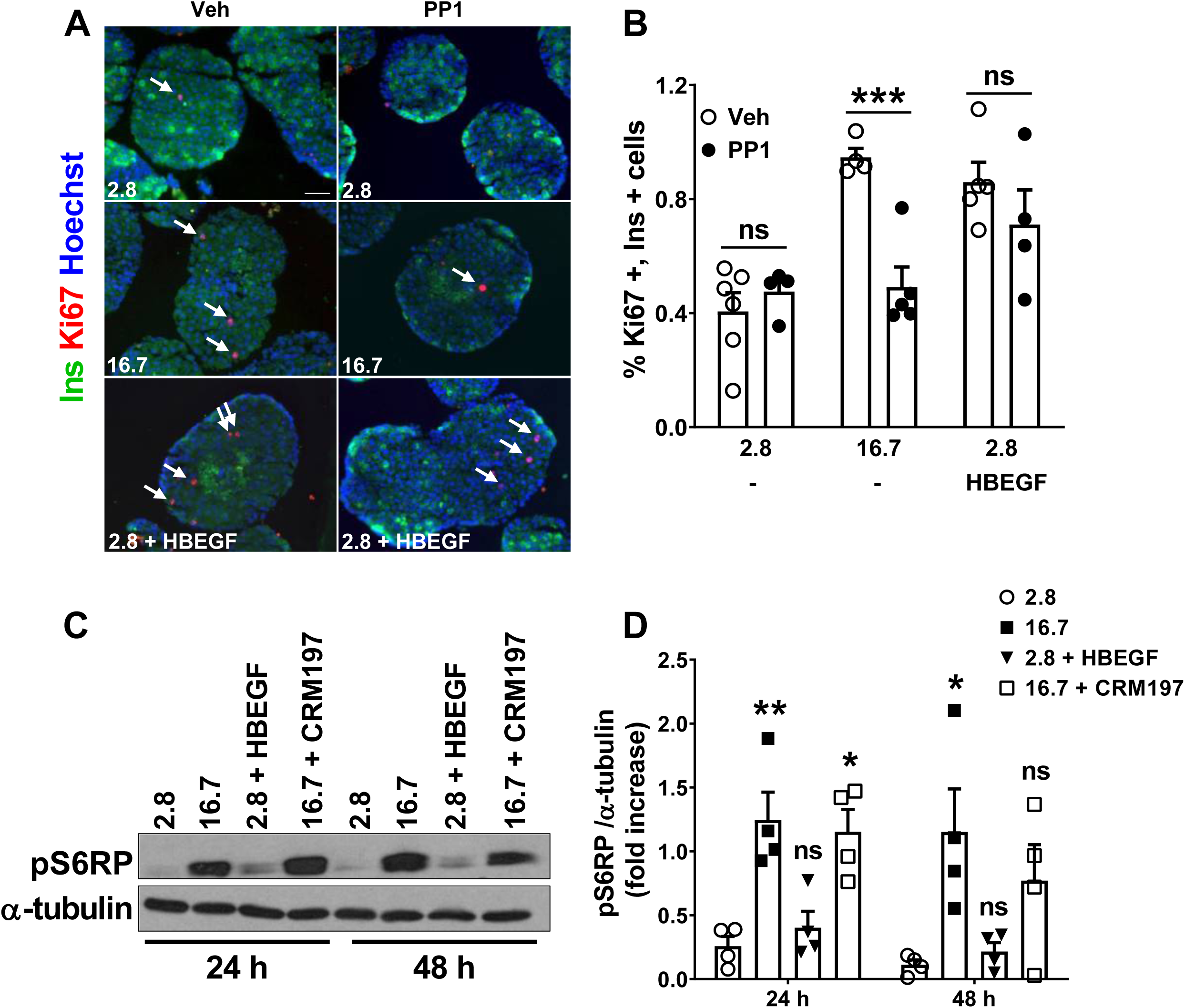
Src is required for glucose-but not HB-EGF-induced β-cell proliferation and glucose-induced mTOR activation does not require HB-EGF. (A, B) Isolated rat islets were exposed to 2.8 or 16.7 mM glucose or HB-EGF (100 ng/ml) in the presence of 2.8 mM glucose for 72 h with or without the Src inhibitor PP1 (1 μM). Proliferation was assessed by Ki67 staining and insulin (Ins). (A) Representative images of insulin (green), Ki67 (red) and nuclei (blue). Arrows show positive nuclei for Ki67. (B) The percentage of Ki67+ insulin+ cells of total insulin+ cells. Scale bars, 50 µm. (C, D) Isolated rat islets were exposed to 2.8 or 16.7 mM glucose or HB-EGF (100 ng/ml) in the presence of 2.8 mM glucose with or without CRM197 (10 μg/ml) for 24 and 48 h. Representative Western blot (C) of phospho-S6RP (pS6RP) and α-tubulin and densitometric quantification (D) of pS6RP normalized to α-tubulin. Data represent individual values and means ± SEM (n=4-6). *p < 0.05, **p<0.01, ***p<0.001, as compared to the 2.8 mM glucose condition or as indicated in the graph (B). ns, not significant.

Glucose-induced β-cell proliferation is dependent on mTOR activation (16). As the mitogenic effect of HB-EGF was also dependent on mTOR in rat islets (Fig. 1G&H), we asked whether mTOR activation by glucose is dependent on HB-EGF. Exposing islets for 24 and 48 h to 16.7 mM glucose led to a significant increase in phosphorylation of the mTOR substrate S6 ribosomal protein (S6RP) (Fig. 7C&D). However, HB-EGF did not increase S6RP phosphorylation, and blocking HB-EGF with CRM197 did not affect glucose-induced mTOR activation (Fig. 7C&D). Hence, glucose activation of mTOR is independent of HB-EGF.

## DISCUSSION

The results of this study demonstrate a critical role for HB-EGF in glucose-induced β-cell proliferation in rat β cells. Exposing isolated islets to exogenous HB-EGF induced β-cell proliferation, whereas blocking HB-EGF signaling by inhibiting either EGFR or HB-EGF completely prevented the proliferative response. *In vivo*, silencing HB-EGF prevented the increase in β-cell proliferation in islets transplanted under the kidney capsule of glucose-infused rats. Taken together, our data identify a glucose/HB-EGF/EGFR axis that controls β-cell proliferation. Mechanistically, we showed that HB-EGF gene expression is induced by glucose in the β cell through the action of ChREBP. In addition we found that glucose-, but not HB-EGF-induced β-cell proliferation is blocked by Src inhibition. As Src family kinases are involved in EGFR transactivation via ADAM metalloproteases, we propose a mechanism whereby glucose activates ChREBP and Src to promote HB-EGF gene expression and HB-EGF membrane shedding, respectively, and subsequently EGFR downstream signaling and cell cycle activation (Fig. 8).

**Figure 8.**
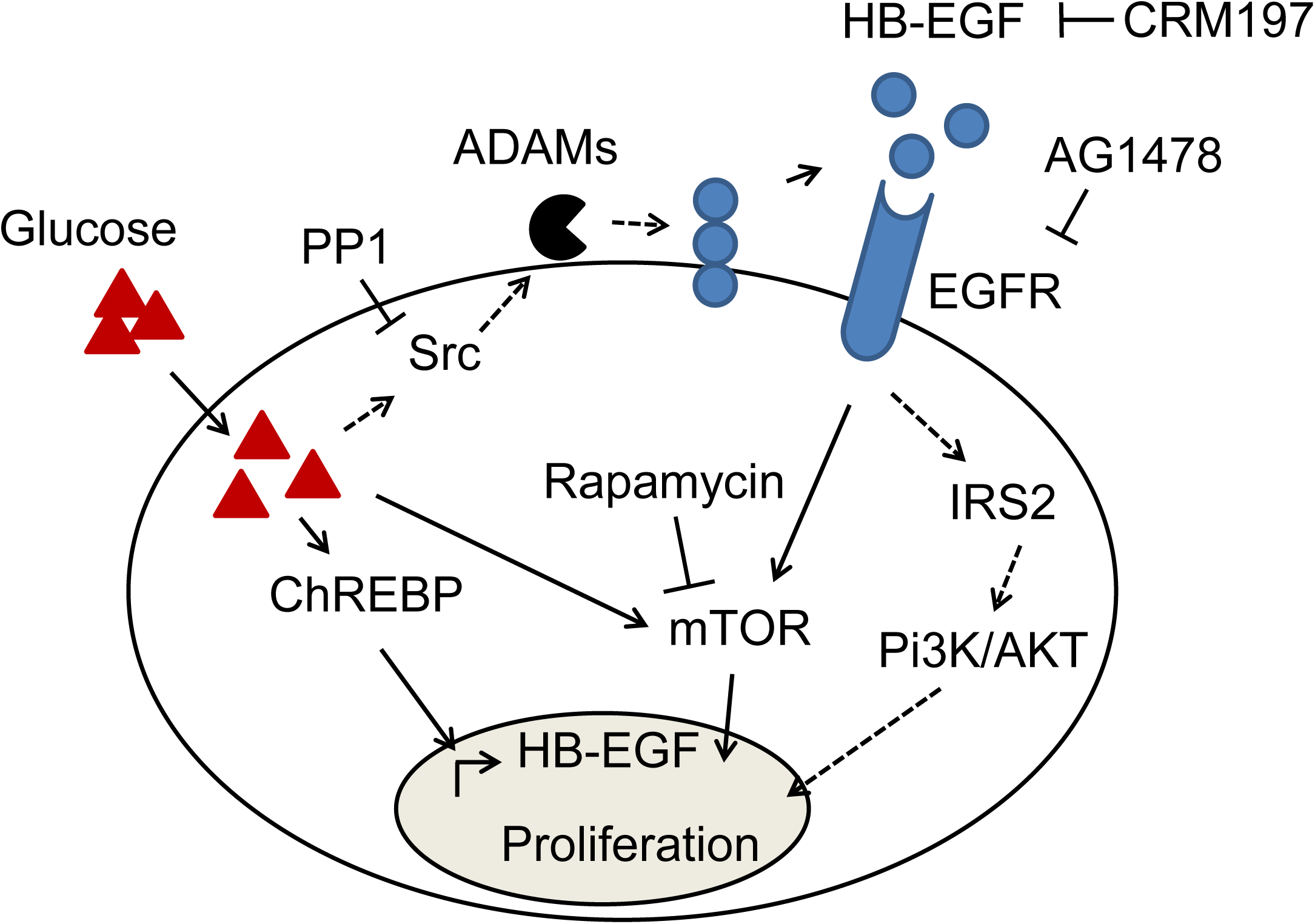
Proposed mechanism of glucose/HB-EGF/EGFR axis controlling β-cell proliferation. An increase in the soluble, active form of HB-EGF is mediated by glucose-induced ChREBP, which increases HB-EGF gene expression, and by glucose-induced Src, which is coupled to metalloprotease (ADAM)-dependent proHB-EGF processing. Subsequent binding of HB-EGF to the β-cell EGFR activates signaling pathways including mTOR but also possibly MAPK, PI3K/AKT and IRS2 that together promote β-cell proliferation. Specific inhibitors used in this study to block glucose- and HB-EGF-induced β-cell proliferation are indicated.

Our previous (21) and current results are in agreement with studies showing that overexpression of HB-EGF by retrograde injection of adenoviruses into the pancreatic duct leads to proliferation of pre-existing β cells in adult mice (31). In contrast, no increase in β-cell proliferation was observed following HB-EGF expression in developing mouse β cells (32). However, the presence of pancreatic fibrosis, stromal expansion and islet dysfunction in this model may have precluded such an effect. Interestingly, overexpression of HB-EGF (31) or BTC (33) in pancreatic ducts promotes β-cell neogenesis, and EGF gain-of-function studies in human duct cells (34; 35) support a similar conclusion. Hence, we propose that the major effect of HB-EGF is to promote proliferation of existing β cells, but that β-cell neogenesis could also contribute to its overall beneficial effects on β-cell mass. In contrast to its effects on β-cell proliferation, acute and extended (24 h) exposure to HB-EGF did not alter insulin secretion or insulin content in rat islets *ex vivo*. However, positive, anti-diabetic, effects of HB-EGF on the β cell were demonstrated in multiple low-dose streptozotocin diabetic mice whereby combined treatment of gastrin and HB-EGF led to improved islet function due in part to a reduction in insulitis (36). Further studies will be required to fully elucidate the pleotropic effects of HB-EGF on pancreatic islets.

We found that AG1478, a specific inhibitor of EGFR with minimal activity towards other ErbB isoforms, completely abrogates HB-EGF induced β-cell proliferation. As HB-EGF signals via EGFR (ErbB1) and ErbB4 but not ErbB2 or ErbB3 (37) and EGFR is expressed in β-cells, whereas ErbB4 is only weakly expressed in rodent islets (38), we propose that HB-EGF acts predominantly via EGFR to promote β-cell proliferation.

EGFR inhibition, loss-of-function and dominant negative studies in adult rodents in the context of pathophysiological and physiological metabolic stress (6; 21; 39), and partial pancreatectomy (22) suggest that β-cell EGFR underlies the maintenance of glucose homeostasis by transducing signals that increase β-cell proliferation and mass. Notwithstanding a role for BTC downstream of GLP-1 (40), however, attempts to investigate the role of EGFR ligands in the regulation of β-cell mass and function have been limited to gain-of-function approaches (31; 33; 41; 42), whereas the identification of endogenous ligands contributing to β-cell compensation is unknown. We found that glucose, a key effector of regulation of β-cell mass in the face of increased insulin demand (11), requires HB-EGF signaling. When rat islets were exposed to glucose *ex vivo* or *in vivo*, the β-cell mitogenic response was dependent on both EGFR and HB-EGF. Although HB-EGF was essential for the glucose response, whether HB-EGF is the sole endogenous EGFR ligand acting during β-cell compensation to metabolic stress remains an open question. BTC (41), epiregulin (EPGN) (43), TGFalpha and EGF (41; 44) exert mitogenic effects on the β cell and are expressed in developing (45) and adult (38) rodent islets and during β-cell neogenesis (46). Hence, different EGFR ligands likely contribute to β-cell compensation in a context-dependent manner.

In previous studies we showed that HB-EGF gene expression is upregulated in islets following nutrient infusion in rats (21) and a similar trend was found in obese, diabetes-resistant (B6) mice (47). Our present results suggest that the increase in HB-EGF gene expression is due, at least in part, to the direct action of glucose. They are consistent with the time- and dose-dependent increase in HB-EGF gene expression observed in response to glucose in INS-1 cells (28) and rat islets (48). In addition we found that ChREBP is necessary for HB-EGF gene expression and that ChREBP binds a 3’ HB-EGF gene enhancer element. Primary targets of ChREBP in the β-cell include RORg and Myc (28), whereas the cell cycle regulatory cyclins and cyclin dependent kinases (cdks), which lack ChREBP binding sites, respond to glucose in a delayed manner due to their dependency on first-phase factors (17; 28). Hence, downstream of ChREBP, HB-EGF/EGFR signaling could play a role alongside first-phase transcription factors to drive cell cycle regulators and initiate β-cell cycle progression in response to glucose (Fig. 8).

Although membrane-anchored proHB-EGF may be involved in juxtacrine signaling (49), the major effects of HB-EGF in the β cell are likely mediated by the soluble form generated by proteolytic processing of proHB-EGF. In mesangial cells, glucose promotes HB-EGF shedding and EGFR transactivation through Src-dependent activation of metalloproteases (30). Our results showing that Src inhibition blocked glucose-but not HB-EGF-induced β-cell proliferation suggest that this phenomenon is also operative in β cells. Consistent with this possibility, short-term exposure of MIN6 and human islets to glucose leads to phosphorylation of the Src family kinase YES (50). Overall, our data are consistent with the model proposed in Fig. 8 whereby glucose promotes Src-dependent proHB-EGF processing leading to HB-EGF shedding and stimulation of β-cell proliferation via paracrine and/or autocrine signaling through the EGFR.

mTOR is an essential mediator of mitogen induced β-cell proliferation (51). Blocking mTOR activity prevents the mitogenic effects of glucose (16) and, as we showed in the present study, also mitigates HB-EGF-induced proliferation. Surprisingly however, blocking HB-EGF had no effect on the increase in mTOR activity in response to glucose yet HB-EGF inhibition completely prevented glucose-induced β-cell proliferation. Hence, we postulate the existence of a parallel signal emanating from EGFR acting alongside the mTOR pathway that is necessary for β-cell cycle engagement. A number of signaling effectors are known to act downstream of EGFR including, MAPK, PI3K/AKT (21) and IRS2 (52) that could contribute to the mitogenic response to HB-EGF (Fig. 8). Full characterization of the signaling pathway linking EGFR to the β-cell mitogenic response will require additional studies examining the potential implication of these kinases.

In conclusion, this study reveals a critical role of HB-EGF/EGFR signaling in glucose-induced β-cell proliferation in rat islets. Future studies will focus on further elucidating the underlying mechanism and assessing the importance of this pathway in human islet pathophysiology.

## ABBREVIATIONS

3C: Chromosome conformation capture
BTC: betacellulin
ChIP: Chromatin immunoprecipitation
ChREBP: Carbohydrate response element binding protein
EGF: Epidermal growth factor
EGFR: Epidermal growth factor receptor
FACS: Fluorescent activated cell sorting
GIR: Glucose infusion rate
HB-EGF: Heparin-binding epidermal growth factor-like growth factor
mTORC1: Mammalian target of rapamycin complex 1
TSS: Transcriptional start site

## ACKNOWLEDGMENTS

This study was supported by the National Institutes of Health (grant R01-DK-58096 to V.P. and R01-DK-108905 to D.K.S.) and the Canadian Institutes of Health Research (grant MOP 77686 to V.P.). H.M. was supported by a doctoral studentship from the Fonds de Recherche Québec - Santé. V.P. holds the Canada Research Chair in Diabetes and Pancreatic Beta Cell Function.

We thank A. Levert (CRCHUM) for technical assistance with isolated experiments and R. Screaton (Sunnybrook Research Institute, Toronto, ON, Canada) for advice on human islet culture.

H.M. and M.R.M. designed the experiments and acquired the data. H.M., M.R.M., D.K.S., J.G. and V.P. researched data, analyzed the results, and wrote the manuscript. All authors revised the manuscript and approved the final version. V.P. is the guarantor of this work and, as such, takes full responsibility for the work.

The authors have no relevant conflict of interest to disclose.

Parts of this study were presented at the 77^th^ Scientific Sessions of the American Diabetes association, San Diego, CA, 9-13 June 2017 and at the 79^th^ Scientific Sessions of the American Diabetes association, San Francisco, CA, 7-11 June 2019.

## REFERENCES

1. Kahn SE, Cooper ME, Del Prato S: Pathophysiology and treatment of type 2 diabetes: perspectives on the past, present, and future. Lancet 2014;383:1068–1083

2. Dor Y, Brown J, Martinez OI, Melton DA: Adult pancreatic beta-cells are formed by self-duplication rather than stem-cell differentiation. Nature 2004;429:41–46

3. Teta M, Rankin MM, Long SY, Stein GM, Kushner JA: Growth and regeneration of adult beta cells does not involve specialized progenitors. Dev Cell 2007;12:817–826

4. Okada T, Liew CW, Hu J, Hinault C, Michael MD, Krtzfeldt J, Yin C, Holzenberger M, Stoffel M, Kulkarni RN: Insulin receptors in beta-cells are critical for islet compensatory growth response to insulin resistance. Proc Natl Acad Sci U S A 2007;104:8977–8982

5. Kim H, Toyofuku Y, Lynn FC, Chak E, Uchida T, Mizukami H, Fujitani Y, Kawamori R, Miyatsuka T, Kosaka Y, Yang K, Honig G, van der Hart M, Kishimoto N, Wang J, Yagihashi S, Tecott LH, Watada H, German MS: Serotonin regulates pancreatic beta cell mass during pregnancy. Nat Med 2010;16:804–808

6. Hakonen E, Ustinov J, Mathijs I, Palgi J, Bouwens L, Miettinen PJ, Otonkoski T: Epidermal growth factor (EGF)-receptor signalling is needed for murine beta cell mass expansion in response to high-fat diet and pregnancy but not after pancreatic duct ligation. Diabetologia 2011;54:1735–1743

7. El Ouaamari A, Dirice E, Gedeon N, Hu J, Zhou JY, Shirakawa J, Hou L, Goodman J, Karampelias C, Qiang G, Boucher J, Martinez R, Gritsenko MA, De Jesus DF, Kahraman S, Bhatt S, Smith RD, Beer HD, Jungtrakoon P, Gong Y, Goldfine AB, Liew CW, Doria A, Andersson O, Qian WJ, Remold-O’Donnell E, Kulkarni RN: SerpinB1 Promotes Pancreatic beta Cell Proliferation. Cell Metab 2016;23:194–205

8. Moulle VS, Vivot K, Tremblay C, Zarrouki B, Ghislain J, Poitout V: Glucose and fatty acids synergistically and reversibly promote beta cell proliferation in rats. Diabetologia 2017;60:879–888

9. Alonso LC, Yokoe T, Zhang P, Scott DK, Kim SK, O’Donnell CP, Garcia-Ocana A: Glucose infusion in mice: a new model to induce beta-cell replication. Diabetes 2007;56:1792–1801

10. Terauchi Y, Takamoto I, Kubota N, Matsui J, Suzuki R, Komeda K, Hara A, Toyoda Y, Miwa I, Aizawa S, Tsutsumi S, Tsubamoto Y, Hashimoto S, Eto K, Nakamura A, Noda M, Tobe K, Aburatani H, Nagai R, Kadowaki T: Glucokinase and IRS-2 are required for compensatory beta cell hyperplasia in response to high-fat diet-induced insulin resistance. J Clin Invest 2007;117:246–257

11. Porat S, Weinberg-Corem N, Tornovsky-Babaey S, Schyr-Ben-Haroush R, Hija A, Stolovich-Rain M, Dadon D, Granot Z, Ben-Hur V, White P, Girard CA, Karni R, Kaestner KH, Ashcroft FM, Magnuson MA, Saada A, Grimsby J, Glaser B, Dor Y: Control of pancreatic beta cell regeneration by glucose metabolism. Cell Metab 2011;13:440–449

12. Stamateris RE, Sharma RB, Hollern DA, Alonso LC: Adaptive beta-cell proliferation increases early in high-fat feeding in mice, concurrent with metabolic changes, with induction of islet cyclin D2 expression. Am J Physiol Endocrinol Metab 2013;305:E149–159

13. Levitt HE, Cyphert TJ, Pascoe JL, Hollern DA, Abraham N, Lundell RJ, Rosa T, Romano LC, Zou B, O’Donnell CP, Stewart AF, Garcia-Ocana A, Alonso LC: Glucose stimulates human beta cell replication in vivo in islets transplanted into NOD-severe combined immunodeficiency (SCID) mice. Diabetologia 2011;54:572–582

14. Martinez SC, Cras-Méneur C, Bernal-Mizrachi E, Permutt MA: Glucose Regulates Foxo1 Through Insulin Receptor Signaling in the Pancreatic Islet β-cell. Diabetes 2006;55:1581–1591

15. Kulkarni RN, Bruning JC, Winnay JN, Postic C, Magnuson MA, Kahn CR: Tissue-specific knockout of the insulin receptor in pancreatic beta cells creates an insulin secretory defect similar to that in type 2 diabetes. Cell 1999;96:329–339

16. Stamateris RE, Sharma RB, Kong Y, Ebrahimpour P, Panday D, Ranganath P, Zou B, Levitt H, Parambil NA, O’Donnell CP, Garcia-Ocana A, Alonso LC: Glucose Induces Mouse beta-Cell Proliferation via IRS2, MTOR, and Cyclin D2 but Not the Insulin Receptor. Diabetes 2016;65:981–995

17. Metukuri MR, Zhang P, Basantani MK, Chin C, Stamateris RE, Alonso LC, Takane KK, Gramignoli R, Strom SC, O’Doherty RM, Stewart AF, Vasavada RC, Garcia-Ocana A, Scott DK: ChREBP mediates glucose-stimulated pancreatic beta-cell proliferation. Diabetes 2012;61:2004–2015

18. Zhang P, Kumar A, Katz LS, Li L, Paulynice M, Herman MA, Scott DK: Induction of the ChREBPbeta Isoform Is Essential for Glucose-Stimulated beta-Cell Proliferation. Diabetes 2015;64:4158–4170

19. Stoeckman AK, Ma L, Towle HC: Mlx is the functional heteromeric partner of the carbohydrate response element-binding protein in glucose regulation of lipogenic enzyme genes. J Biol Chem 2004;279:15662–15669

20. Fontes G, Zarrouki B, Hagman DK, Latour MG, Semache M, Roskens V, Moore PC, Prentki M, Rhodes CJ, Jetton TL, Poitout V: Glucolipotoxicity age-dependently impairs beta cell function in rats despite a marked increase in beta cell mass. Diabetologia 2010;53:2369–2379

21. Zarrouki B, Benterki I, Fontes G, Peyot ML, Seda O, Prentki M, Poitout V: Epidermal growth factor receptor signaling promotes pancreatic beta-cell proliferation in response to nutrient excess in rats through mTOR and FOXM1. Diabetes 2014;63:982–993

22. Song Z, Fusco J, Zimmerman R, Fischbach S, Chen C, Ricks DM, Prasadan K, Shiota C, Xiao X, Gittes GK: Epidermal Growth Factor Receptor Signaling Regulates beta Cell Proliferation in Adult Mice. J Biol Chem 2016;291:22630–22637

23. Taylor SR, Markesbery MG, Harding PA: Heparin-binding epidermal growth factor-like growth factor (HB-EGF) and proteolytic processing by a disintegrin and metalloproteinases (ADAM): a regulator of several pathways. Semin Cell Dev Biol 2014;28:22–30

24. Kelpe CL, Johnson LM, Poitout V: Increasing triglyceride synthesis inhibits glucose-induced insulin secretion in isolated rat islets of langerhans: a study using adenoviral expression of diacylglycerol acyltransferase. Endocrinology 2002;143:3326–3332

25. Ferdaoussi M, Bergeron V, Zarrouki B, Kolic J, Cantley J, Fielitz J, Olson EN, Prentki M, Biden T, Macdonald PE, Poitout V: G protein-coupled receptor (GPR)40-dependent potentiation of insulin secretion in mouse islets is mediated by protein kinase D1. Diabetologia 2012;55:2682–2692

26. Hagman DK, Latour MG, Chakrabarti SK, Fontes G, Amyot J, Tremblay C, Semache M, Lausier JA, Roskens V, Mirmira RG, Jetton TL, Poitout V: Cyclical and alternating infusions of glucose and intralipid in rats inhibit insulin gene expression and Pdx-1 binding in islets. Diabetes 2008;57:424–431

27. Ghislain J, Fontes G, Tremblay C, Kebede MA, Poitout V: Dual-Reporter beta-Cell-Specific Male Transgenic Rats for the Analysis of beta-Cell Functional Mass and Enrichment by Flow Cytometry. Endocrinology 2016;157:1299–1306

28. Schmidt SF, Madsen JG, Frafjord KO, Poulsen L, Salo S, Boergesen M, Loft A, Larsen BD, Madsen MS, Holst JJ, Maechler P, Dalgaard LT, Mandrup S: Integrative Genomics Outlines a Biphasic Glucose Response and a ChREBP-RORgamma Axis Regulating Proliferation in beta Cells. Cell Rep 2016;16:2359–2372

29. Hagege H, Klous P, Braem C, Splinter E, Dekker J, Cathala G, de Laat W, Forne T: Quantitative analysis of chromosome conformation capture assays (3C-qPCR). Nat Protoc 2007;2:1722–1733

30. Taniguchi K, Xia L, Goldberg HJ, Lee KW, Shah A, Stavar L, Masson EA, Momen A, Shikatani EA, John R, Husain M, Fantus IG: Inhibition of Src kinase blocks high glucose-induced EGFR transactivation and collagen synthesis in mesangial cells and prevents diabetic nephropathy in mice. Diabetes 2013;62:3874–3886

31. Kozawa J, Tokui Y, Moriwaki M, Li M, Ohmoto H, Yuan M, Zhang J, Iwahashi H, Imagawa A, Yamagata K, Tochino Y, Shimomura I, Higashiyama S, Miyagawa J: Regenerative and therapeutic effects of heparin-binding epidermal growth factor-like growth factor on diabetes by gene transduction through retrograde pancreatic duct injection of adenovirus vector. Pancreas 2005;31:32–42

32. Means AL, Ray KC, Singh AB, Washington MK, Whitehead RH, Harris RC, Jr., Wright CV, Coffey RJ, Jr., Leach SD: Overexpression of heparin-binding EGF-like growth factor in mouse pancreas results in fibrosis and epithelial metaplasia. Gastroenterology 2003;124:1020–1036

33. Tokui Y, Kozawa J, Yamagata K, Zhang J, Ohmoto H, Tochino Y, Okita K, Iwahashi H, Namba M, Shimomura I, Miyagawa J: Neogenesis and proliferation of beta-cells induced by human betacellulin gene transduction via retrograde pancreatic duct injection of an adenovirus vector. Biochem Biophys Res Commun 2006;350:987–993

34. Rescan C, Le Bras S, Lefebvre VH, Frandsen U, Klein T, Foschi M, Pipeleers DG, Scharfmann R, Madsen OD, Heimberg H: EGF-induced proliferation of adult human pancreatic duct cells is mediated by the MEK/ERK cascade. Lab Invest 2005;85:65–74

35. Suarez-Pinzon WL, Lakey JR, Brand SJ, Rabinovitch A: Combination therapy with epidermal growth factor and gastrin induces neogenesis of human islet {beta}-cells from pancreatic duct cells and an increase in functional {beta}-cell mass. J Clin Endocrinol Metab 2005;90:3401–3409

36. Castillo GM, Nishimoto-Ashfield A, Banerjee AA, Landolfi JA, Lyubimov AV, Bolotin EM: Omeprazole and PGC-formulated heparin binding epidermal growth factor normalizes fasting blood glucose and suppresses insulitis in multiple low dose streptozotocin diabetes model. Pharm Res 2013;30:2843–2854

37. Elenius K, Paul S, Allison G, Sun J, Klagsbrun M: Activation of HER4 by heparin-binding EGF-like growth factor stimulates chemotaxis but not proliferation. EMBO J 1997;16:1268–1278

38. DiGruccio MR, Mawla AM, Donaldson CJ, Noguchi GM, Vaughan J, Cowing-Zitron C, van der Meulen T, Huising MO: Comprehensive alpha, beta and delta cell transcriptomes reveal that ghrelin selectively activates delta cells and promotes somatostatin release from pancreatic islets. Mol Metab 2016;5:449–458

39. Hakonen E, Ustinov J, Palgi J, Miettinen PJ, Otonkoski T: EGFR signaling promotes beta-cell proliferation and survivin expression during pregnancy. PLoS One 2014;9:e93651

40. Buteau J, Foisy S, Joly E, Prentki M: Glucagon-like peptide 1 induces pancreatic beta-cell proliferation via transactivation of the epidermal growth factor receptor. Diabetes 2003;52:124–132

41. Huotari MA, Palgi J, Otonkoski T: Growth factor-mediated proliferation and differentiation of insulin-producing INS-1 and RINm5F cells: identification of betacellulin as a novel beta-cell mitogen. Endocrinology 1998;139:1494–1499

42. Li L, Seno M, Yamada H, Kojima I: Promotion of beta-cell regeneration by betacellulin in ninety percent-pancreatectomized rats. Endocrinology 2001;142:5379–5385

43. Kuntz E, Broca C, Komurasaki T, Kaltenbacher MC, Gross R, Pinget M, Damge C: Effect of epiregulin on pancreatic beta cell growth and insulin secretion. Growth Factors 2005;23:285–293

44. Krakowski ML, Kritzik MR, Jones EM, Krahl T, Lee J, Arnush M, Gu D, Mroczkowski B, Sarvetnick N: Transgenic expression of epidermal growth factor and keratinocyte growth factor in beta-cells results in substantial morphological changes. J Endocrinol 1999;162:167–175

45. Huotari MA, Miettinen PJ, Palgi J, Koivisto T, Ustinov J, Harari D, Yarden Y, Otonkoski T: ErbB signaling regulates lineage determination of developing pancreatic islet cells in embryonic organ culture. Endocrinology 2002;143:4437–4446

46. Li M, Miyagawa J, Moriwaki M, Yuan M, Yang Q, Kozawa J, Yamamoto K, Imagawa A, Iwahashi H, Tochino Y, Yamagata K, Matsuzawa Y: Analysis of expression profiles of islet-associated transcription and growth factors during beta-cell neogenesis from duct cells in partially duct-ligated mice. Pancreas 2003;27:345–355

47. Tu Z, Keller MP, Zhang C, Rabaglia ME, Greenawalt DM, Yang X, Wang IM, Dai H, Bruss MD, Lum PY, Zhou YP, Kemp DM, Kendziorski C, Yandell BS, Attie AD, Schadt EE, Zhu J: Integrative analysis of a cross-loci regulation network identifies App as a gene regulating insulin secretion from pancreatic islets. PLoS Genet 2012;8:e1003107

48. Bensellam M, Van Lommel L, Overbergh L, Schuit FC, Jonas JC: Cluster analysis of rat pancreatic islet gene mRNA levels after culture in low-, intermediate- and high-glucose concentrations. Diabetologia 2009;52:463–476

49. Ray KC, Blaine SA, Washington MK, Braun AH, Singh AB, Harris RC, Harding PA, Coffey RJ, Means AL: Transmembrane and soluble isoforms of heparin-binding epidermal growth factor-like growth factor regulate distinct processes in the pancreas. Gastroenterology 2009;137:1785–1794

50. Yoder SM, Dineen SL, Wang Z, Thurmond DC: YES, a Src family kinase, is a proximal glucose-specific activator of cell division cycle control protein 42 (Cdc42) in pancreatic islet beta cells. J Biol Chem 2014;289:11476–11487

51. Balcazar N, Sathyamurthy A, Elghazi L, Gould A, Weiss A, Shiojima I, Walsh K, Bernal-Mizrachi E: mTORC1 activation regulates beta-cell mass and proliferation by modulation of cyclin D2 synthesis and stability. J Biol Chem 2009;284:7832–7842

52. Oh YS, Shin S, Lee YJ, Kim EH, Jun HS: Betacellulin-induced beta cell proliferation and regeneration is mediated by activation of ErbB-1 and ErbB-2 receptors. PLoS One 2011;6:e23894

