## Supplemental Material for "HB-EGF Signaling is Required for Glucose-Induced Pancreatic β-Cell Proliferation in Rats"

**Supplemental Table S1:** Sources of antibodies used.

| Antibody | Dilution/Concentration | Company | Cat # |
|---|---|---|---|
| **Immunohistochemistry** | | | |
| Insulin | 1:500 | DAKO | A0564 |
| Nkx6.1 | 5 ug/ml | DSHB | F55A12 |
| Ki67 | 1:400 | Abcam | 15580 |
| PHH3 | 1:600 | Novus | NB21-1091 |
| **Immunoblotting** | | | |
| p-S6 ribosomal protein | 1:2500 | Cell Signaling | 2211S |
| alpha-tubulin | 1:20000 | Abcam | 4074 |
| **Immunoprecipitation** | | | |
| ChREBP | - | Abcam | ab157153 |
| Normal IgG | - | Santa Cruz | sc-2027 |

**Supplemental Table S2.** 3C, ChIP and RT-PCR primers.

| Primer name | forward | Reverse or anchor |
|---|---|---|
| **3C** | | |
| 2 3C | TCCCAAGCCACGTATGATGT | GGAGACACGGGCTCTTACCT |
| 3 3C | GTGTTGGGAATGGGATTTTG | GGTAAGAGCCCGTGTCTCCG |
| 4 3C | AACACGAGGAAGGGAGGAAG | GGTAAGAGCCCGTGTCTCCG |
| 5 3C | GACAACTGGGGGACTGAGAC | GGTAAGAGCCCGTGTCTCCG |
| 6 3C | AAGCCTGAAATGAACAGAACAA | GGTAAGAGCCCGTGTCTCCG |
| 7 3C | AGGTCCTAGGTGTCCTGGAA | GGTAAGAGCCCGTGTCTCCG |
| 8 3C | TGCATCCCAAGCAACACTAA | GGTAAGAGCCCGTGTCTCCG |
| 17 3C | GAGGGGTTGTGGGTTCATTA | GGTAAGAGCCCGTGTCTCCG |
| 18 3C | AGGTTCTTTCTCCGTTGTGC | GGTAAGAGCCCGTGTCTCCG |
| 19 3C | GTTGGTGACCGGTGAGAGTC | GGTAAGAGCCCGTGTCTCCG |
| 20 3C | GGACCGACTCTGAACAGACA | GGTAAGAGCCCGTGTCTCCG |
| 21 3C | ACATCCAACTTGCCTTCTGC | GGTAAGAGCCCGTGTCTCCG |
| **ChIP** | | |
| Pklr coding region | GTGGAGCACGGTGGTATCTT | CTTCACGCCTTCATGGTTCT |
| HB-EGF downstream | GTGAGTGCCCATGTCACTGT | GCTTGCCTTCCTTCCTTTTT |
| **RT-PCR** | | |
| HB-EGF-1 | GACCGATCTGGACCTTTTCA | CCGTGGATGCAGTAGTCCTT |
| HB-EGF-2 | GTGGGTAGCAGCTGGTTTGT | GTTGGTGACCGGTGAGAGT |
| Cyclophilin A | GCCATTATGGCGTGTGAAGTC | CTTGCTGCAGACATGGTCAAC |

### Figure S1

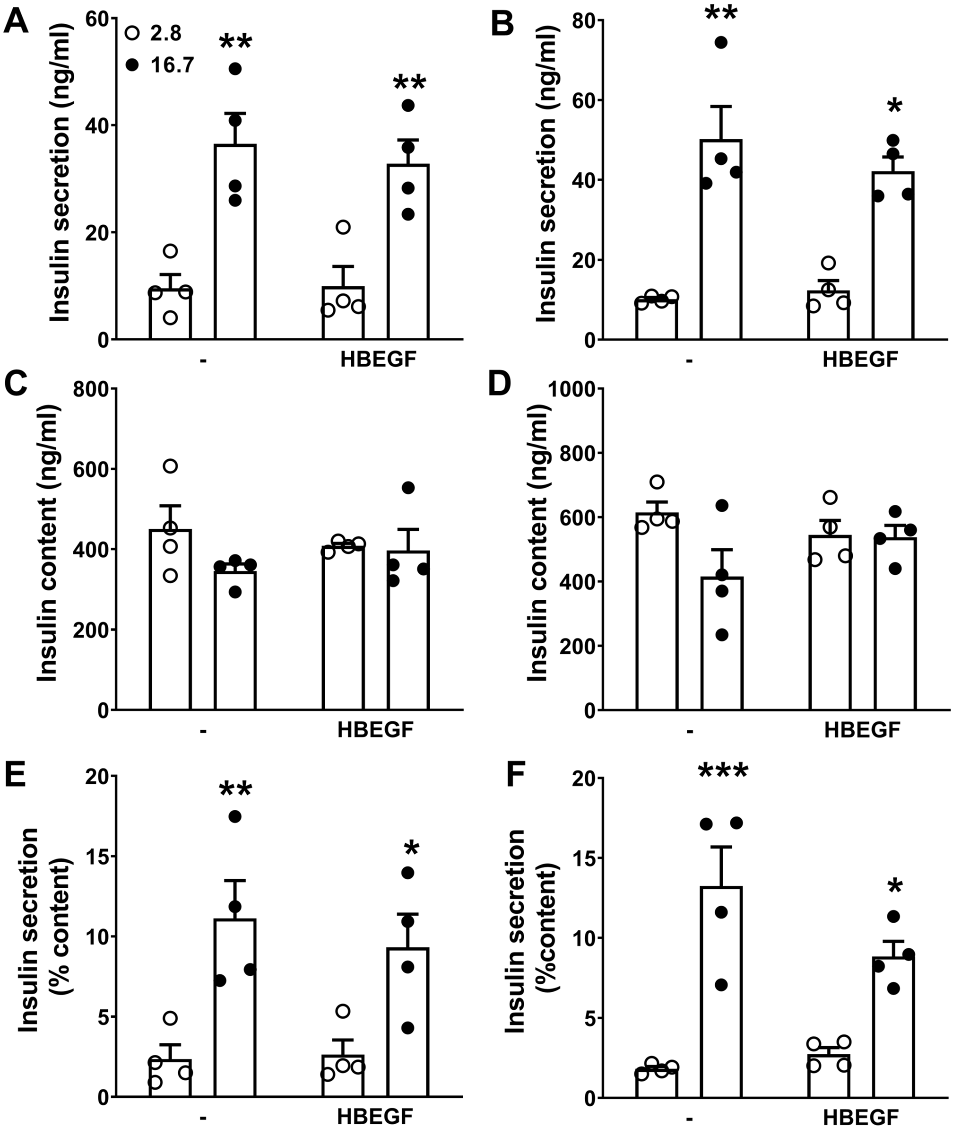


**Figure S1. HB-EGF does not affect glucose-induced insulin secretion or insulin content in rat islets ex vivo.** (A-D) Insulin secretion (A, B) and total islet insulin content (C, D) were determined in 1-h static incubations in response to 2.8 and 16.7 mM glucose in isolated rat islets with or without exposure to HB-EGF (100 ng/ml) during the static incubation with glucose (A, C, E) or 24 h prior (B, D, F). (E, F) Insulin levels are expressed as a percentage of total islet insulin content. Data represent individual values and means ± SEM (n=4). *p < 0.05, **p<0.01, ***p<0.001, as compared to the 2.8 mM glucose condition.

### Figure S2

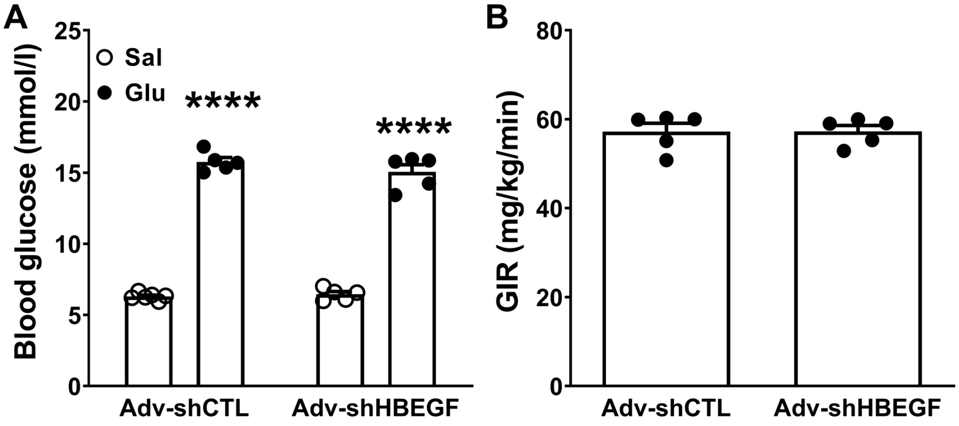


**Figure S2. Blood glucose levels and GIR following glucose infusion in Lewis rats after islet transplant.** Control shRNA adenovirus (shCTL) or shHB-EGF adenovirus (shHBEGF) infected islets were transplanted under the kidney capsule of Lewis rats and infused with saline (Sal) or glucose (Glu) for 72 h. Blood glucose levels (A) and GIR (B) during the infusion. Data represent individual values and means ± SEM (n=4-6). ****$p<0.0001$, as compared to the saline condition.

### Figure S3

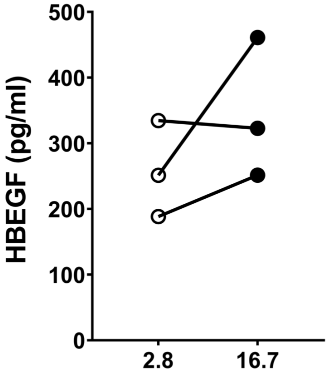


**Figure S3. HB-EGF levels in rat islet extracts.** Isolated rat islets were exposed to 2.8 and 16.7 mM glucose for 1h. HB-EGF was assessed in the protein extract by ELISA kit. Data represent individual values (n=3).
